# ORYX-MRSI: A Fully-Automated Open-Source Software for Three-Dimensional Proton Magnetic Resonance Spectroscopic Imaging Data Analysis

**DOI:** 10.1101/2021.11.12.468398

**Authors:** Sevim Cengiz, Muhammed Yildirim, Abdullah Bas, Esin Ozturk-Isik

**Affiliations:** Institute of Biomedical Engineering, Bogazici University, Istanbul, Turkey; Computer Vision, Mohamed bin Zayed University of Artificial Intelligence, Abu Dhabi, UAE

**Keywords:** Magnetic resonance spectroscopy, MRSI, open-source, software

## Abstract

Proton magnetic resonance spectroscopic imaging (^1^H-MRSI) provides noninvasive evaluation of brain metabolism. However, there are some limitations of 1H-MRSI preventing its wider use in the clinics, including the spectral quality issues, partial volume effect and chemical shift artifact. Additionally, it is necessary to create metabolite maps for analyzing spectral data along with other MRI modalities. In this study, a MATLAB-based open-source data analysis software for 3D ^1^H-MRSI, called Oryx-MRSI, which includes modules for visualization of raw ^1^H-MRSI data and LCModel outputs, chemical shift correction, tissue fraction calculation, metabolite map production, and registration onto standard MNI152 brain atlas while providing automatic spectral quality control, is presented. Oryx-MRSI implements region of interest analysis at brain parcellations defined on MNI152 brain atlas. All generated metabolite maps are stored in NIfTI format. Oryx-MRSI is publicly available at https://github.com/sevimcengiz/Oryx-MRSI along with six example datasets.

## Introduction

Three-dimensional (3D) magnetic resonance spectroscopic imaging (^1^H-MRSI) provides metabolic information about the brain, including its cellularity, myelin degeneration, neuronal loss and dysfunction, and energy storage (Chang et al., 2013; Govindaraju et al., 2000; Miller, 1991). ^1^H-MRSI has been an important tool for the diagnosis, follow-up, and treatment planning of several diseases, such as brain tumors and neurological disorders. (Kantarci et al., 2000; Nelson, 2003, 2011) Despite the vast amount of information provided by ^1^H-MRSI, it is still not widely employed in the clinical settings. As a result, there has been a major effort for improving the clinical utility of ^1^H-MRSI with recent developments in data acquisition, processing, and quantitative analysis aspects (Bartnik-Olson et al., 2021; Chiew et al., 2018; Hingerl et al., 2020; Landheer et al., 2020; Near et al., 2021; Oeltzschner et al., 2019; Oz et al., 2020; Povazan et al., 2020; Tapper et al., 2021; Wilson et al., 2019). As part of these extensive efforts, open-source command-line scripts or software with user-friendly graphical user interfaces (GUIs) have been released in the past few year (Clarke et al., 2021; Crane et al., 2013; Edden et al., 2014; Maudsley et al., 2006, 2009; Naressi et al., 2001; Oeltzschner et al., 2020; Poullet et al., 2007; Provencher, 1993; Reynolds et al., 2006; Simpson et al., 2017; Soher et al., 2011; Wilson et al., 2011; Wilson, 2021). LCModel is one of the most popular MRS data quantification tools, which estimates metabolite concentration and metabolite to total creatine ratios for a range of metabolites, including macromolecules and lipids, and it recently became open source (Provencher, 1993). On the other hand, jMRUI (Naressi et al., 2001) and Tarquin(Reynolds et al., 2006; Wilson et al., 2011) offer customizable GUI-based tools for spectral visualization and quantification. MIDAS (Metabolite Imaging and Data Analysis System) provides whole brain MRSI data visualization, processing, and analysis. Additionally, Osprey is a new open-source MRS data analysis software that currently supports single-voxel MRS data analysis(Oeltzschner et al., 2020). Moreover, FSL-MRS is another Python-based open-source tool that provides data quantification of single-voxel MRS and 2D MRSI after converting the data into the NIfTI format (Clarke et al., 2021). More recently, MRspant has been released, which is an automated R-based MR spectroscopic data analysis tool for reading, visualizing, and processing MRS data (Wilson, 2021).

In this paper, we present an open-source 3D MRSI data analysis software with a user-friendly GUI, named Oryx-MRSI, which reads LCModel outputs as well as raw spectral data and enables visualization and metabolite map generation considering the chemical shift correction while providing automated spectral quality control based on full width at half maximum (FWHM), signal-to-noise ratio (SNR), Cramer–Rao lower bounds (CRLB), and CSF fraction (fCSF). Oryx-MRSI also provides registration of metabolite maps onto the MNI152 brain atlas (Mazziotta et al., 1995) and enables region of interest (ROI) analysis at multiple brain locations, including the functional parcellations of the human cerebral cortex based on resting-state functional MRI (rs-fMRI) networks (Schaefer et al., 2018) or the MNI structural brain atlas regions. The tool exports spreadsheets that include the mean, median, and standard deviation (SD) values of the metabolites and metabolite ratios with or without fCSF correction for multiple brain regions.

## Methods

Oryx-MRSI was written in MATLAB 2020a (Mathworks Inc., Natick, MA, USA) and it has been tested using MATLAB versions 2020a and newer in Ubuntu 18.04.5 LTS and macOS 11.4 Big Sur. All sub-functions of Oryx-MRSI can be called as command-line scripts or Oryx-MRSI can be easily used through a user-friendly GUI. A complete Oryx-MRSI data analysis pipeline includes eight different modules, which are load data, co-registration, segmentation, FWHM and SNR, spectral quality, metabolite maps, registration, and ROI analysis. Before the analysis starts, the user is asked to provide some parameters, including the cut-off values of the CRLB, fCSF, FWHM, and SNR for voxel exclusion criteria, radiofrequency (RF) bandwidth of the sequence for chemical shift correction, and cut-off value for the probabilistic binary map after registration onto the MNI152 brain atlas. The following sections describe the Oryx-MRSI modules in detail.

### Load Data

This module enables the user to visualize 3D ^1^H-MRSI data located either in a raw data file or COORD file outputted by LCModel. Currently, Oryx-MRSI supports 3D ^1^H-MRSI raw data saved in the SPAR/SDAT format acquired on a Philips MR scanner. Some examples of 3D ^1^H-MRSI datasets can be found in the “~/Oryx-MRSI/Dataset” folder under the GitHub repository located at https://github.com/sevimcengiz/Oryx-MRSI. Oryx-MRSI also allows the user to load and visualize their own dataset stored as an SPAR file under the “~/Oryx-MRSI/Dataset” folder. The necessary steps of the data preparation before data analysis with Oryx-MRSI are detailed in the documentation available in the GitHub repository.

The default imaging system for the data order of the raw data and LCModel outputs are left, posterior, and superior (LPS). Accordingly, the column numbers increase from right to left, and the row numbers increase from the anterior to posterior directions at a selected slice at the visualization screen. Although several metabolites, including lipids and macromolecules, are quantified by LCModel, Oryx-MRSI currently creates metabolite maps for total creatine (Cr+PCr), glutamate glutamine complex (Glu+Gln), total choline (GPC+PCh), myoinositol (Ins), lactate (Lac), and lipids (Lip13a, Lip13b, Lip13a+Lip13b). The software also allows zooming in on a voxel for a closer view, and visualization of the individual metabolites.

### Co-registration

In this module, first, the position and orientation information of the scanner-space coordinates of the field of view (FOV) are parsed from an SPAR file, and each voxel’s size, position, and orientation information are calculated considering their slice, row, and column numbers. This module uses ‘Gannetmask_Philips’ function from Gannet for co-registration (Edden et al., 2014) after some necessary modifications for 3D data analysis and generates binary masks for FOV, volume of interest (VOI), and individual voxels. Oryx-MRSI asks the user to select one reference metabolite from among H2O, NAA, Cr, Cho, and Lac/Lip, and to specify the RF bandwidths of the MR system for the excitation and the first and second echo directions in Hz. The chemical shift correction is applied when the chemical shift correction option is set to “on”, and the RF bandwidths are provided by the user.

For chemical shift correction, the gradient strengths (T/mm) on the excitation, and first and second echo directions (dir) are calculated as follows:

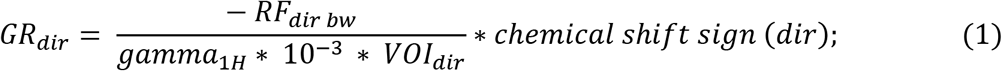

where gamma_1H_ is equal to the gyromagnetic ratio in Hz/T, VOI represents the volume of interest box sizes in mm in the respective directions, chemical shift sign is either +1 or −1, and positive chemical shift directions are L, P, and S (H) for the LPS imaging system. The chemical shift amounts in mm in the three respective directions are then calculated as,

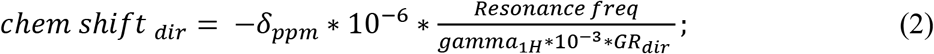

where δ_ppm_ is the ppm difference between the shifted and reference metabolites, and the resonance frequency is in Hz. Currently, Cr+PCr (3.03 ppm), Glu+Gln (2.25 ppm), GPC+PCh (3.2 ppm), Ins (3.52 ppm), Lac (1.3 ppm), Lip13a (1.3 ppm), Lip13b (1.3 ppm), and Lip13a+Lip13b (1.3 ppm) metabolite maps are estimated. The corresponding FOV for these metabolites are shifted in space by their respective chemical shift amounts. This module ensures that the binary voxel masks are positioned onto the same coordinate system and co-registered to the reference structural image. As a result, the resultant binary masks are saved in the NIFTI format under the “~/coreg_binary_mask” folder.

### Segmentation

The segmentation module uses FMRIB’s Software Library (FSL)-Fast (Zhang et al., 2001) tool to segment the T1w-MRI into cerebrospinal fluid (CSF), white matter (WM), and gray matter (GM) regions. If the anatomical reference image for ^1^H-MRSI is T1w-MRI, this module calculates the CSF, WM, and GM fractions at each voxel of all different binary masks, which are FOV placements of every metabolite after chemical shift correction. On the other hand, if the anatomical reference image for ^1^H-MRSI is T2w-MRI, the T1w-MRI and CSF, WM, and GM probabilistic maps are first registered to T2w-MRI using FSL-Flirt (Jenkinson & Smith, 2001). Then, the CSF, WM, and GM fractions are calculated for all voxels of different metabolite masks. The tissue fraction calculations of the 3D ^1^H-MRSI are modified from Osprey (Oeltzschner et al., 2020), which calculates GM, WM, and CSF fractions for single-voxel MRS data. Each metabolite has a different box placement if the chemical shift correction is set to ‘on’. As a result, the tissue fractions are calculated separately for each metabolite in FOV.

### FWHM & SNR

The FWHM & SNR module reads the LCModel TABLE files of multivoxel ^1^H-MRSI data to retrieve the full width at half maximum (FWHM) and signal-to-noise ratio (SNR) information for each voxel. This module also provides a visualization of the FWHM and SNR maps for all slices.

### Automatic Spectral Quality Control

This module provides automatic spectral quality control to select high-quality voxels for the analysis based on the FWHM, SNR, CRLB, and fCSF thresholds provided by the user. The GUI asks the user to determine the cut-off values for FWHM, SNR, CRLB, and fCSF to exclude poor quality spectra. Each metabolite has a CRLB value provided in TABLE files after LCModel data analysis, indicating the quantification reliability, which were used to exclude spectra based on CRLB.

### Metabolite Maps

All LCModel TABLE files were parsed to obtain the concentration values of the metabolites. These results are positioned onto a 3D MR volume with the same image space and properties as the reference anatomical MRI. The off-center, size, and angulation along the anterior-posterior (ap), left-right (lr), and cranial–caudal (cc) directions were considered to create several 3D MR spectroscopic maps including both the concentration and CSF-corrected concentration maps, and their Ins or Cr + PCr ratio maps. CSF correction is necessary to reduce the partial volume effect in multi-voxel MRSI data analysis (Quadrelli et al., 2016). The corrected metabolite concentrations are calculated as follows:

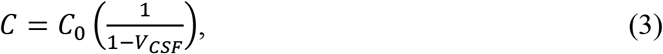

where C represents the corrected metabolite concentration and C_0_ the initial metabolite concentration obtained from the LCModel TABLE files. V_CSF_ is the volume fraction of the CSF of each voxel calculated at the segmentation module. The outputs of this module are saved under the “spectra/nifti” directory.

### Registration

The registration module enables the user to register the reference anatomical MRI onto the MNI152 brain atlas using the FSL-FLIRT tool to obtain a transformation matrix, which is then used to register the spectral image volumes, including the binary mask of the VOI and all the raw and CSF-corrected concentration or ratio maps onto the atlas. Although the original binary mask of the VOI has all the ones inside and zeros outside, the pixel intensities have a range of probabilistic values ranging between 0 and 1 after registration. The user is asked to provide an inclusion cut-off value for probabilistic maps. Thus, only those pixels that exceed this threshold are considered to be within the VOI and considered for further analysis. The outputs of this module are saved under the “~/spectra/nifti/MNI_Regist_Probabilistic” directory.

### ROI Analyze

This module enables the user to evaluate the metabolic maps at functional parcellations of the human cerebral cortex on rs-fMRI networks or MNI structural brain atlas regions and calculates the mean, median, and standard deviation of the chosen concentration map in these brain regions. If the number of pixels in an ROI is greater than the exclusion ratio (given on the ROI module page), then that ROI is included in the analysis of the metabolite of interest, otherwise, it is not. The ROI Analyze module exports all the results in a Microsoft Excel sheet.

### Data Acquisition of Example Datasets

Examples of 3D ^1^H-MRSI datasets for Oryx-MRSI are available in the “~/Oryx-MRSI/Dataset” folder which is under the GitHub repository. One healthy control and one patient with Parkinson’s disease were scanned on a 3T clinical MR scanner (Philips Healthcare, Best, Netherlands) after obtaining written informed consent. The study was conducted with the approval of the Institutional Review Board. The brain MRI protocol included T1w MRI (TR = 8.31 ms, TE = 3.81 ms, flip angle=8, acquisition matrix = 256 × 256 × 90, FOV = 240 mm × 240 mm, slice thickness = 1 mm, scan time = 143 s), T2w MRI (TR = 10243 ms, TE = 80 ms, flip angle = 90 °, acquisition matrix = 128 × 128 × 90, FOV = 240 mm × 240 mm × 180 mm, slice thickness = 2 mm, scan time = 3.5 min), and a 3D ^1^H-MRSI acquired using a point resolved spectroscopy (PRESS) sequence (TR = 1000 ms, TE = 52 ms, 1000 Hz, 1024 points, data acquisition matrix = 14 x 14 x 3, 588 voxels, FOV = 140 mm × 140 mm × 36 mm, voxel size = 10 mm × 10 mm ×12 mm, total scan time = 8 min). A T2w MRI was used as the reference anatomical MR image for ^1^H-MRSI. The excitation, echo, and echo2 directions were along AP, RL, and FH, respectively. The phase-encoding direction (RFOV) was along RL. The chemical shift directions were defined during the data acquisition, which were A, L, and F along AP, LR, and foot-head (FH) for the example datasets, respectively. The reference metabolite was NAA at 2.02 ppm.The raw ^1^H-MRSI data were quantified with LCModel (Provencher, 1993) using a simulated basis set named gamma_press_te52_128mhz_627d.basis provided by the LCModel distributor. Later, brain extracted NIfTI files of T1w MRI, T2w MRI, and the raw data files in SPAR and SDAT format, LCModel outputs, and the screenshots of the ^1^H-MRSI data acquisition were saved under “~/Oryx-MRSI/Dataset/Patient_Name” directory. The spectral datasets given in the GitHub repository were evaluated in detail qualitatively, and the exclusion criteria for the example dataset were defined as <8, >0.10, >30, and >0.30 for the SNR, FWHM, CRLB, and fCSF values, respectively.

MRI brain protocols for Braino GE Phantom trials are given as text files in the GitHub repository, separately. The Braino GE Phantom was scanned on the same scanner with four different trials to assess the chemical shift directions (TR/TE=1000/52 ms, 3D scan mode, transverse orientation). In the first and second trials, the RFOV was set as RL. The chemical shift directions were defined as A, L, and F along AP, LR, and FH, respectively. The only difference between the first and second trials was the plan scan metabolite, which was set as H2O and NAA, respectively. In the third and fourth trials, the RFOV was set as AP. The chemical shift directions were A, R, and H along AP, R, and FH, respectively. The plan scan metabolites of the third and fourth trials were H2O and NAA, respectively.

## Results

Figure 1 shows the main page of Oryx-MRSI, where the user can provide the required parameters of the cut-off values of CRLB, fCSF, FWHM, SNR for voxel exclusion criteria, RF bandwidth of the sequence for chemical shift correction, and cut-off value for the probabilistic binary map after registration onto the MNI152 brain atlas.

**Figure 1.**
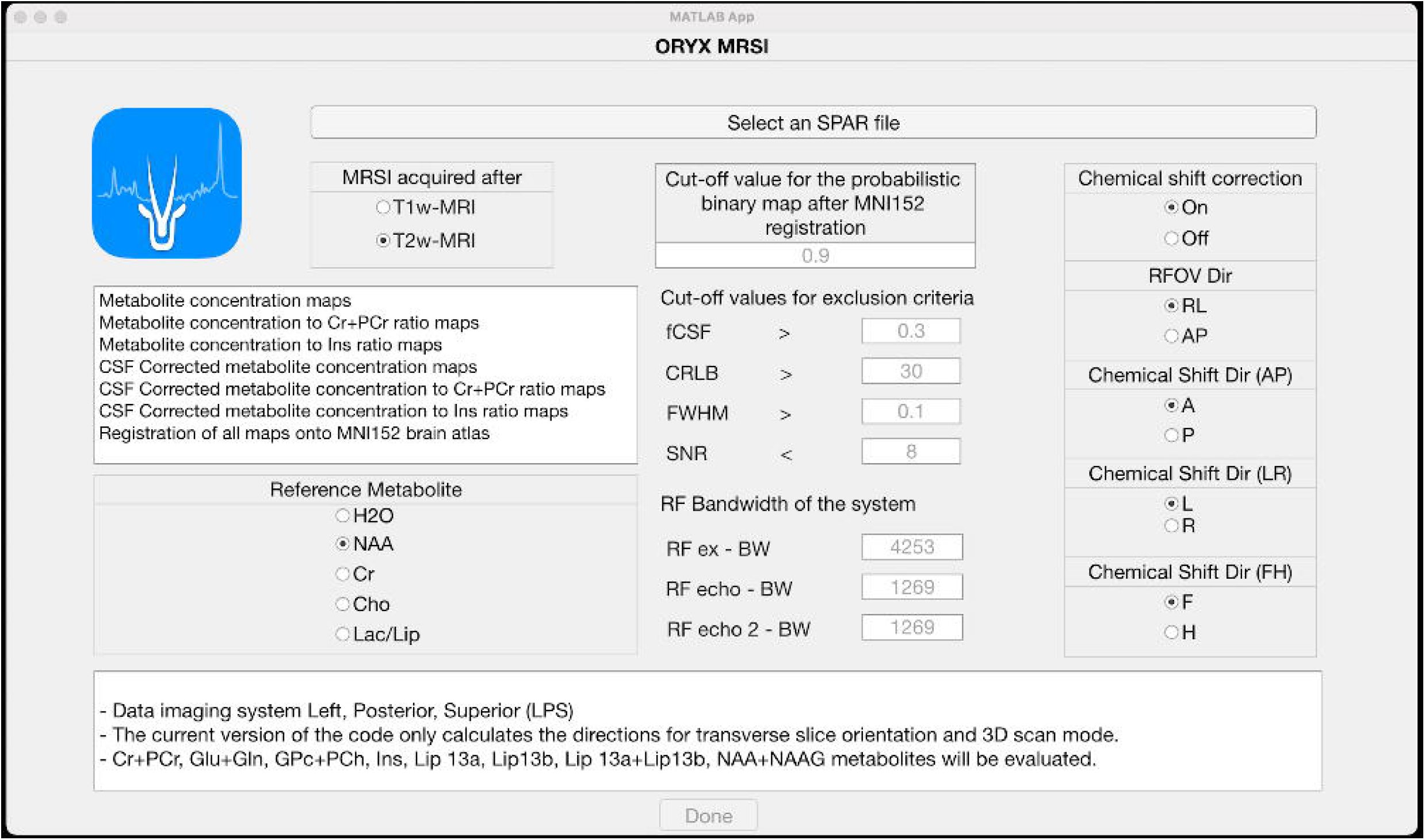
The main page of Oryx-MRSI where the user could provide the required parameters of cut-off values of CRLB, fCSF, FWHM, and SNR for voxel exclusion criteria, RF bandwidth of the sequence for chemical shift correction, and the cut-off value for the probabilistic binary map after registration onto the MNI152 brain atlas

The dataset named K_01 in the GitHub repository was used for the example data analysis. Oryx-MRSI successfully loaded the example dataset and enabled a visualization of the 3D ^1^H-MRSI dataset after reading either the raw data (Figure 2A) or the COORD files (Figure 2B). The software also supported zooming in on a voxel for a closer view (Figure 2C), and visualization of the individual metabolite fits (Figure 2D).

**Figure 2.**
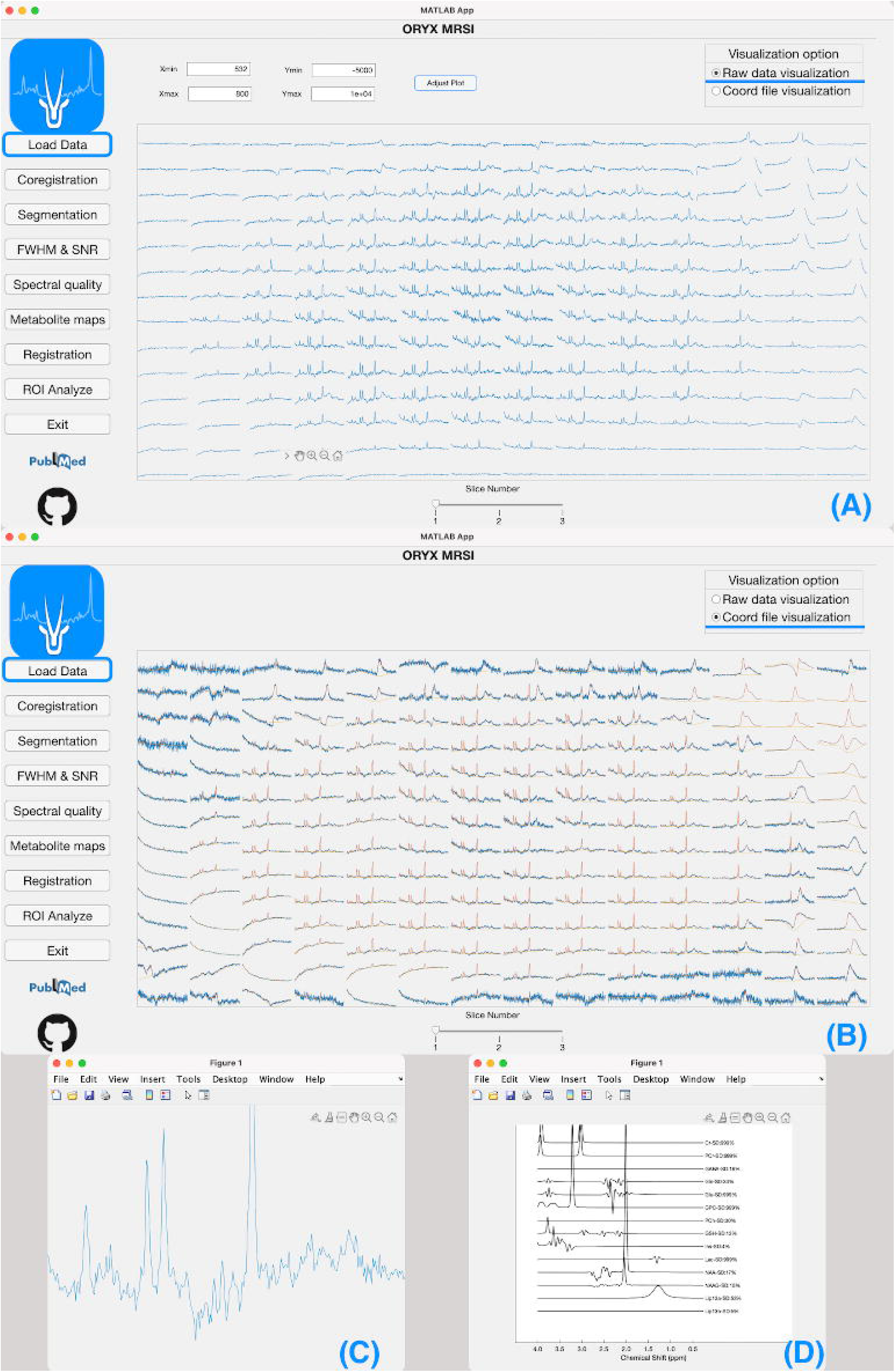
A, Visualization of the 3D ^1^H-MRSI dataset after reading the raw data. B, Visualization of the 3D ^1^H-MRSI dataset after reading the COORD files. C, Zooming in on a voxel for a closer view. D, Visualization of the individual metabolite fits

Figure 3A shows example binary FOV masks of Cr+PCr and Lac placed on an anatomical T2-weighted MRI, which indicates the importance of taking chemical shift into account for ^1^H-MRSI. Figure 3B depicts the FOV and a single voxel (in white, slice =1, row=1, and col=1) out of 3 × 14 × 14 voxels for the NAA+NAAG (blue box), Cr+PCr (green box), and Lac (red box) metabolites.

**Figure 3.**
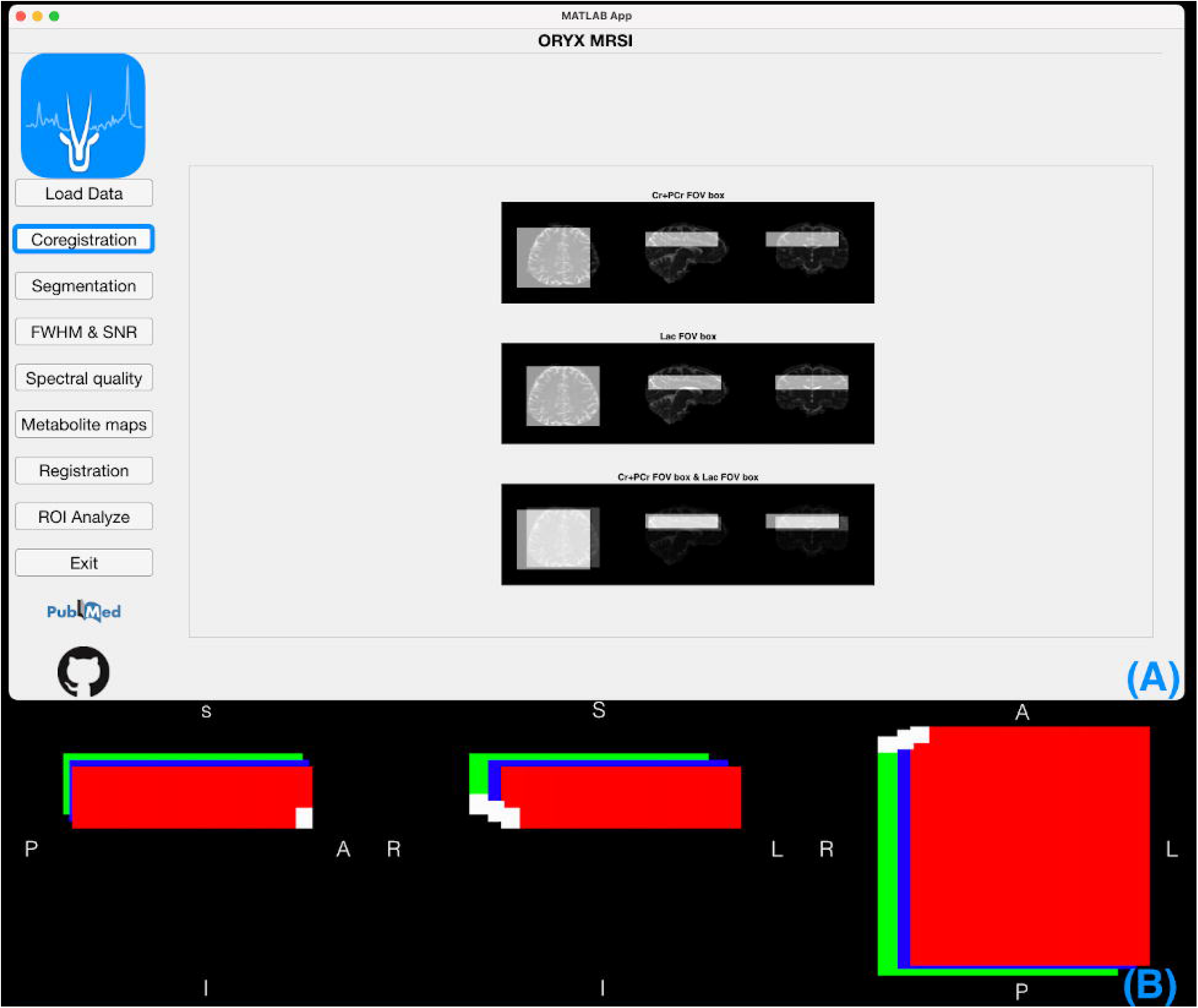
A, Example binary FOV masks of Cr+PCr and Lac placed on an anatomical T2-weighted MRI. B, The FOV and a single voxel (in white, slice =1, row=1, and col=1) out of 3 × 14 × 14 voxels for the NAA+NAAG (blue box), Cr+PCr (green box), and Lac (red box) metabolites

Figure 4 shows the fCSF, fWM, and fGM maps of the NAA+NAAG box at slice 1 of the example dataset (Figure 4A) along with the FWHM, and SNR maps (Figure 4B). and the voxels included in the analysis after the quality check (Figure 4C).

**Figure 4.**
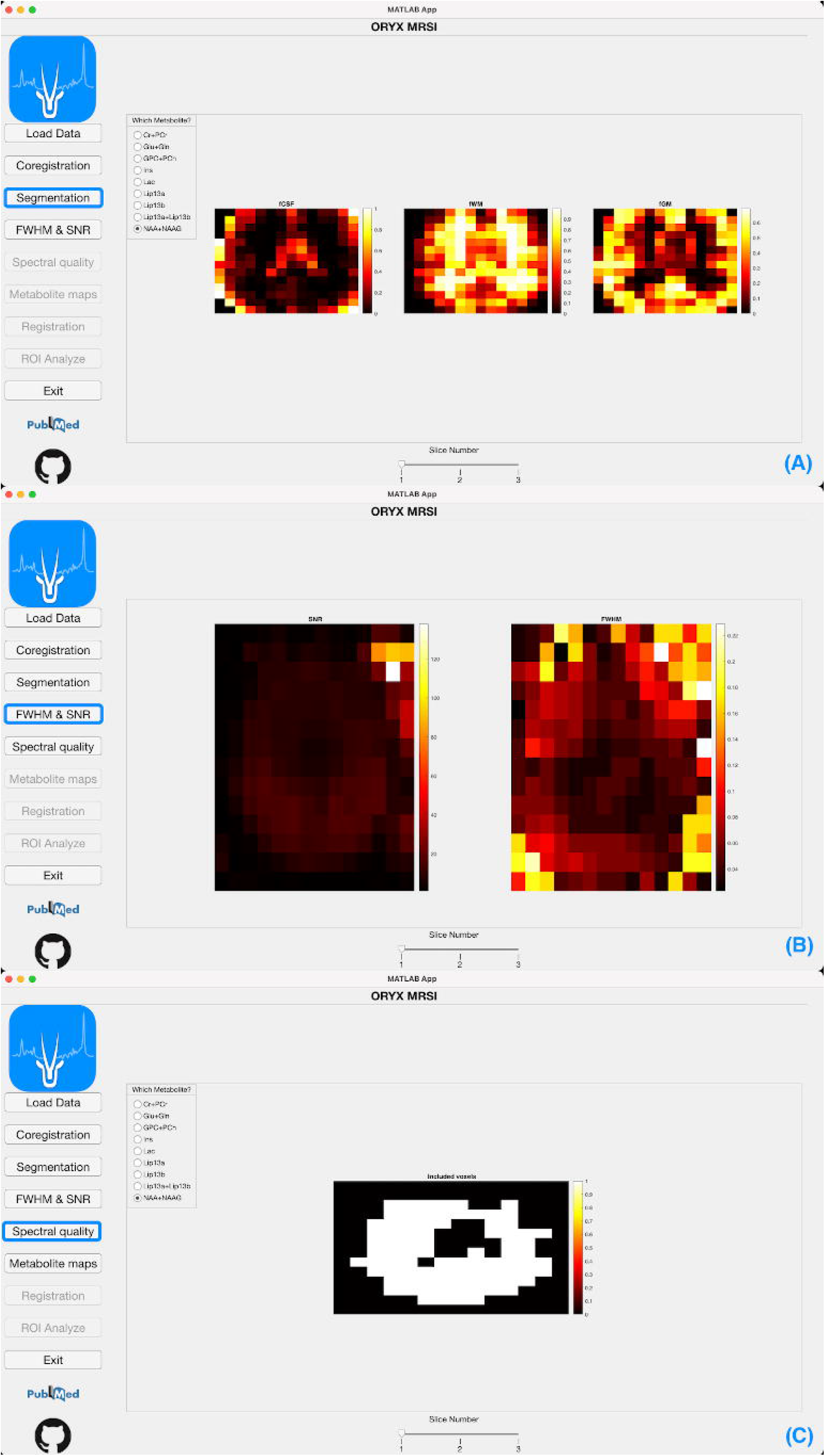
A, The fCSF, fWM, and fGM maps of the NAA+NAAG box at slice 1 of the example dataset. B, The FWHM and SNR maps. C, The voxels included in the analysis after the quality check

The NAA+NAAG concentration map was generated using the metabolite map module (Figure 5A). Additionally, this module allows for the visualization of CSF corrected concentration maps, and metabolite to Ins or Cr+PCr ratio maps. The analysis results revealed that the metabolite to Cr+PCr ratio values estimated by LCModel and Oryx-MRSI did not agree particularly well, because LCModel does not consider the chemical shift while Oryx-MRSI recalculates metabolite concentrations to tCr or mI ratio values at every voxel after chemical shift correction. An NAA+NAAG concentration map after registration onto the MNI152 brain atlas is shown in Figure 5B. Finally, an example spreadsheet file with the statistical results is shown in Figure 6.

**Figure 5.**
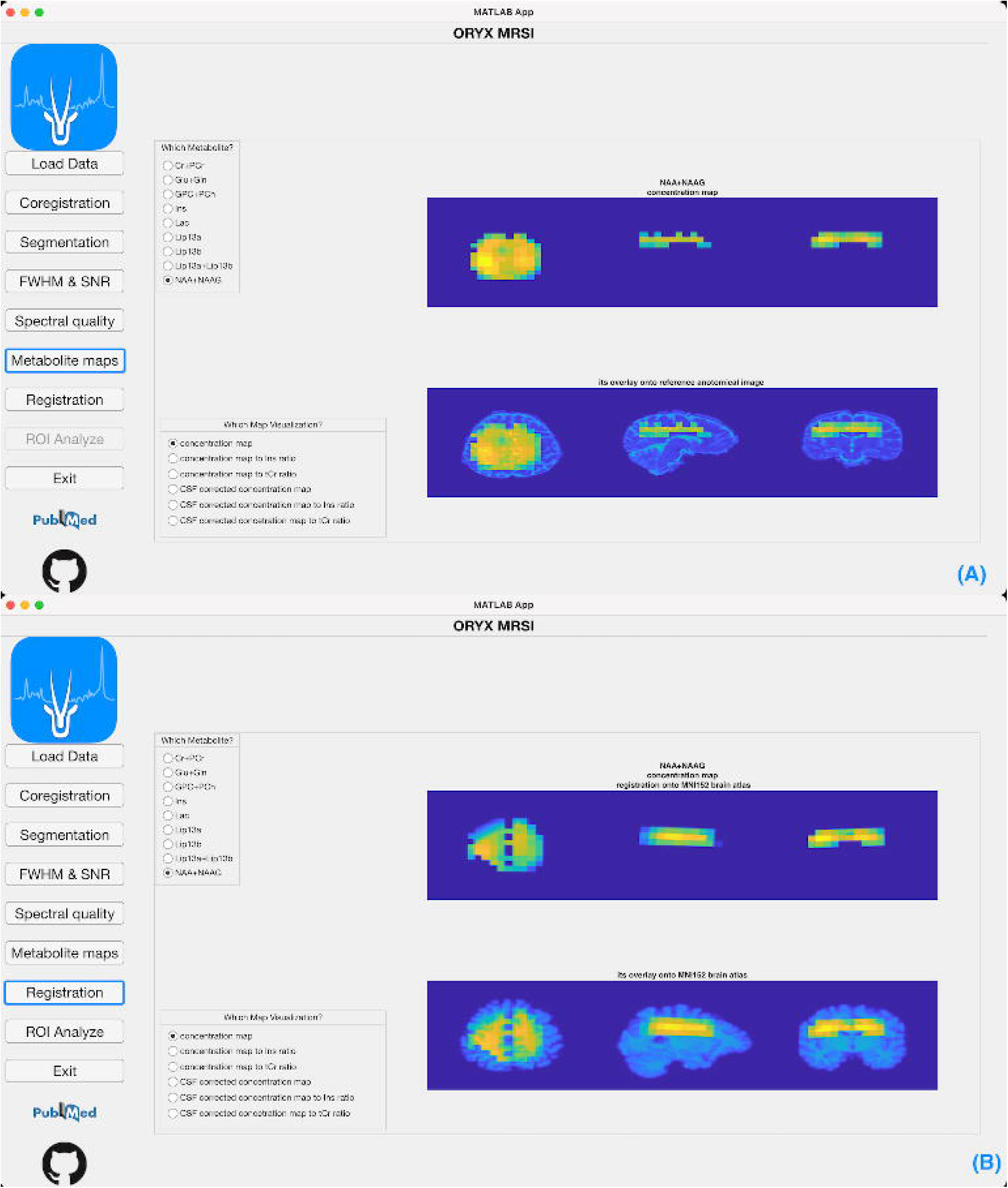
A, The NAA+NAAG concentration map was generated using the metabolite map module. B, An NAA+NAAG concentration map after registration onto the MNI152 brain atlas

**Figure 6.**
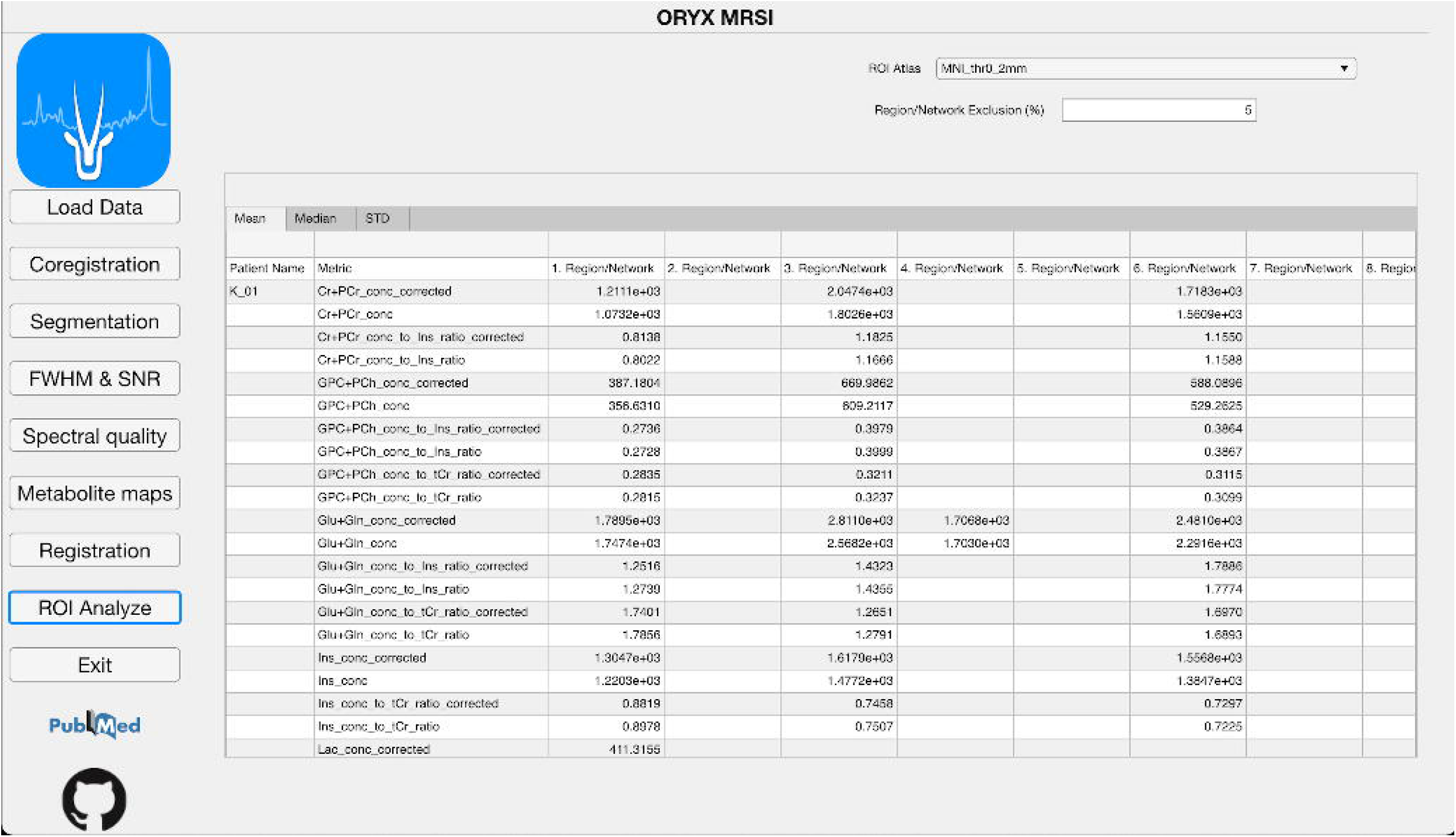
An example spreadsheet file with the statistical results

Figure 7 shows the chemical shift directions of the Cr (green) and Lac (red) boxes when NAA (blue) was set as the reference metabolite on the second phantom trial (Figure 7A, RFOV = RL, chemical shift directions = A, L, F), and fourth phantom trial (Figure 7B, RFOV = AP, chemical shift directions = A, R, H). The Lac box was shifted toward (A, L, F) and (A, L, H) directions, whereas there were shifts toward (P, R, H) and (P, R, F) for Cr in the second and fourth phantom trials, respectively. All the calculated chemical shift directions were consistent with those displayed on the Philips MR scanner console. Two slices of the ^1^H-MRSI data acquired with the first and second phantom trials, which were conducted with water (A) and NAA (B) as the plan scan metabolites, respectively, are shown in Figure 8. All metabolites were shifted toward the left direction when water was set as the reference frequency. However, Cho was shifted toward the right and Lac was shifted toward the left directions when NAA was used as the reference frequency. The chemical shift amount in the AP direction was less than in the other directions due to the higher RF pulse bandwidth.

**Figure 7.**
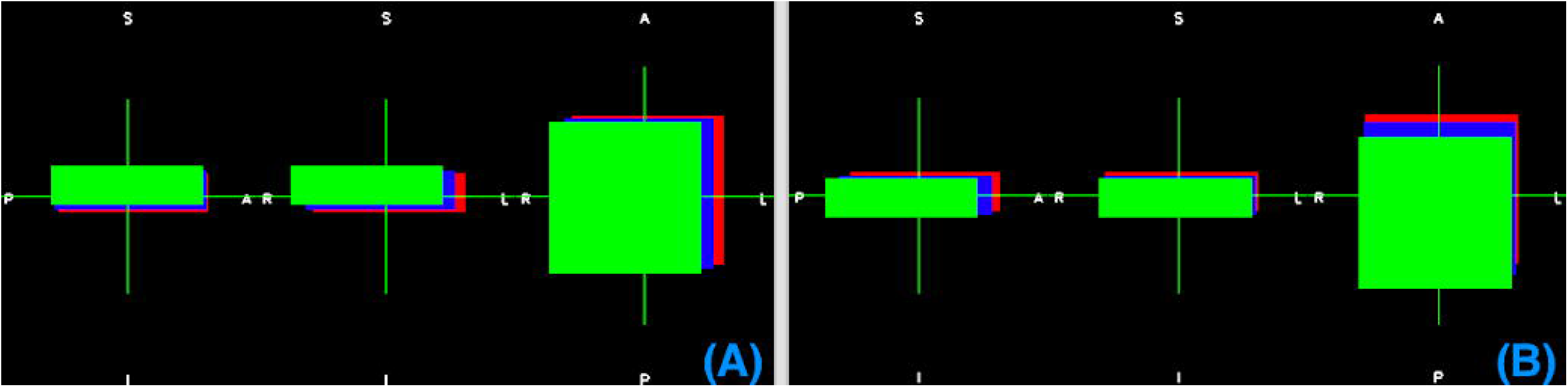
The chemical shift directions of the Cr (green) and Lac (red) boxes when NAA (blue) was set as the reference metabolite on the second phantom trial (Figure 7A, RFOV = RL, chemical shift directions = A, L, F), and fourth phantom trial (Figure 7B, RFOV = AP, chemical shift directions = A, R, H)

**Figure 8.**
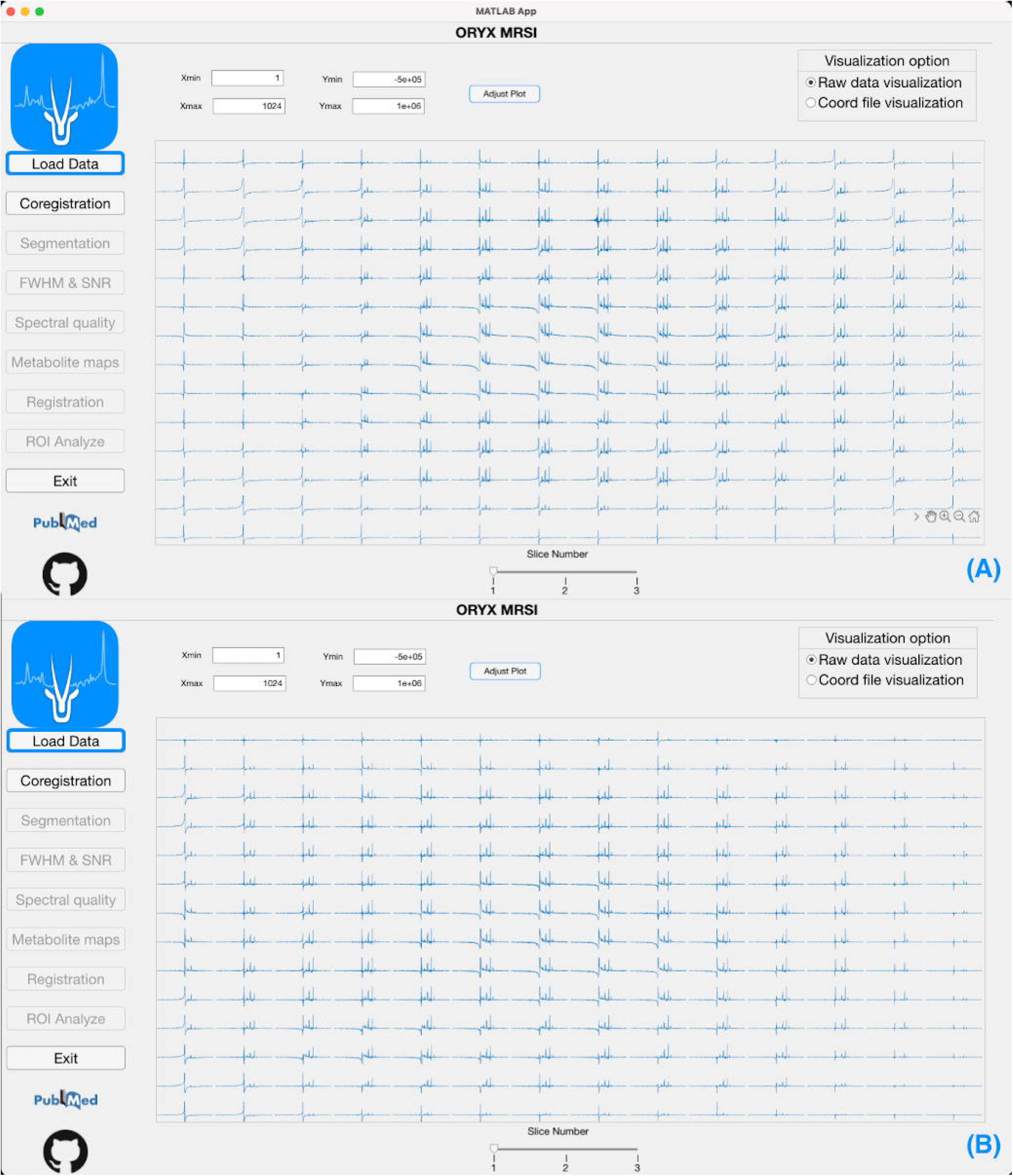
Two slices of the ^1^H-MRSI data acquired with the first and second phantom trials, which were conducted with water (A) and NAA (B) as the plan scan metabolites, respectively

## Discussion

^1^H-MRSI is a clinically useful technique that provides metabolic information useful for the diagnosis, treatment decision making, and follow-up for several diseases including brain disorders (Chang et al., 2013; Kantarci et al., 2000; Nelson, 2003, 2011). There have been several multivoxel ^1^H-MRSI studies that have employed data analysis pipelines including chemical shift correction, tissue fraction calculation, metabolite map generation and registration onto common brain atlases, and ROI analysis (Andronesi et al., 2020; Hangel et al., 2018; Hingerl et al., 2020; Malaspina et al., 2021; Maudsley et al., 2010; Parikh et al., 2015; Solanky et al., 2020). However, a standardized data analysis software has not yet been developed that can execute all these steps for the analysis of 3D ^1^H-MRSI data. This study presents Oryx-MRSI, which is an open-source MATLAB-based end-to-end pipeline for complementary MRSI data analysis after data quantification. Oryx-MRSI supports chemical shift and tissue-fraction corrections and generation of MNI-registered metabolite maps after considering several data quality criteria. Importantly, all metabolite map outputs are stored in a standard medical image file type, NIfTI, which is a common data storage format for neuroimaging that can easily be visualized using FSLEyes (McCarthy, 2020), SPM (Frackowiak et al., 1997), MRICron (Rorden et al., 2007), or NiBabel in Python (Brett et al., 2019). Additionally, Oryx-MRSI generates brain-atlas-based statistical analysis results.

Many studies have reported the importance of CSF correction after anatomical image segmentation (Bonekamp et al., 2005; Degaonkar et al., 2005; Horska et al., 2002; Quadrelli et al., 2016; Tal et al., 2012). While LCModel (Provencher, 1993) does not take into account partial volume effect, Osprey (Oeltzschner et al., 2020), FSL-MRS (Clarke et al., 2021), and MRSpant (Wilson, 2021) provide corrections for it. Similarly, Oryx-MRSI also supports partial volume fraction calculations and CSF correction. Another important factor in ^1^H-MRSI data quantification is the chemical shift effect. (Ozturk-Isik et al., 2006; Wilson et al., 2019) As a result, Oryx-MRSI has a chemical shift correction module. However, it is important to note that chemical shift correction formula is dependent on the specifics of the spectroscopy sequence. Additionally, although the PRESS sequence suffers from this artifact, semi-LASER sequences have better localization performance.

The production of metabolite maps registered to standardized brain atlases, such as the MNI152 brain atlas, is required to facilitate group-based statistical analysis (Bhogal et al., 2020) or to analyze spectroscopic data along with other MR images, such as arterial spin labeling (ASL) MRI (Cengiz, et al., 2017, Cengiz, et al., 2018). FSL-MRS, Osprey, and MRSpant co-register MRS data onto the reference anatomical MRI to compute the CSF fraction and correct for it, but they currently do not support metabolite map generation, registration onto a common brain atlas, and ROI analysis. On the other hand, Oryx-MRSI has these additional features.

Another requirement for reliable data analysis is automated quality control of the spectra based on the linewidth, SNR, and accuracy of the peak fits (Dong et al., 2009). The CRLB is commonly employed to assess the quality of the data quantification. However, Kreis reported that the use of CRLB values to assess the spectral quality might affect the resultant findings (Kreis, 2016); hence, we enabled Oryx-MRSI to assess the effects of different CRLB thresholds on data analysis.

This study had some limitations. It is necessary to note that the chemical shift directions and formulations provided in the Methods section were calculated for a single MR vendor, and it is necessary to validate these formulations for different vendors. Additionally, Oryx-MRSI currently only supports transverse slice orientation and 3D scan mode. Moreover, LCModel data quantification results are currently needed to activate FWHM& SNR, spectral quality, metabolite maps, registration, ROI analyze sections. Oryx-MRSI could be installed and run only on macOS and Linux, because FSL does not directly run on Windows operating systems but requires a Windows Subsystem for Linux (WSL). Additionally, Oryx-MRSI currently supports ROI based analysis, and a voxel-based statistical analysis module will be developed in the future. Oryx-MRSI will be continuously updated to provide support for different MR vendors, and possible integration with earlier open-source MRS data analysis tools.

## Conclusions

Oryx-MRSI is a fully automated open-source software for comprehensive data analysis of 3D ^1^H-MRSI that includes specific modules for automated spectral quality control, metabolite map production, co-registration with anatomical MRI, segmentation of anatomical MRI for CSF fraction correction, registration onto MNI152 brain atlas, and ROI analysis. The metabolic map outputs produced by Oryx-MRSI supports concurrent evaluation of MRSI data along with other MR modalities at brain parcellations defined on MNI152 brain atlas and could enable group-based statistical analysis. As a result, Oryx-MRSI might facilitate more common use of ^1^H-MRSI in clinical settings. This open-source software is publicly available at https://github.com/sevimcengiz/Oryx-MRSI, and a step-by-step data analysis video tutorial is available at https://www.youtube.com/watch?v=X4nqGlny-O8.

## Acknowledgements

This project was funded by the TUBITAK 115S219. We thank all open-source MRI and MRS tools for providing inspiration and some modules that were incorporated into the Oryx-MRSI.

## Ethics approval

All procedures performed in studies involving human participants were in accordance with the ethical standards of the institutional and/or national research committee and with the 1964 Helsinki declaration and its later amendments or comparable ethical standards.

## Consent to participate

Informed consent was obtained from all individual participants included in the study.

## Availability of data and material

The data that support the findings of this study are openly available in https://github.com/sevimcengiz/Oryx-MRSI/tree/main/Dataset.

